# Discriminant Analysis of Principle Component analyses of Physiological Data

**DOI:** 10.1101/2020.01.09.899898

**Authors:** Omar Haidar, Samuel Ball, Richard Barrett-Jolley

## Abstract

There are many situations in physiological and pharmacological analyses where multivariate data is collected. Frequently these are analysed with t-tests and multiple (Bonferroni) comparisons or ANOVA with post-hoc test. Increasingly, even with more powerful computers many variables and it seems that feature reduction would be a useful approach. The most commonly used method is principle component analyses, but in this report we compare this to a technique developed for genetic analyses, discriminant analysis of principle component (DAPC) analyses. A simple to use and well-maintained library exists for DAPC analyses, Adegenet^2^, and using this we find that DAPC detects differences between synthetic physiological datasets with significantly greater accuracy than traditional PCA.

## Introduction

Genomics is inherently a discipline rich in massive datasets; the technology to generate Next Generation Sequencing for example, has come alongside more powerful computers and computation methods to analyse it. Although physiology is a much older discipline, there are often still massive datasets, for example real-time imagine data of Ca^2+^ or other second messengers, and yet these data are typically transformed to a form in which a traditional Student’s t-test or ANOVA can be done (Figure 1). There is relatively little usage of multivariate analyses even if it may be possible, and the tools are freely available as R-packages. The more widespread use of multivariate analyses in physiology has a number of profound advantages, one of which is the ability to analyse entire datasets objectively, without having to, for example, select a region of interest for analysis.

**Figure 1:**
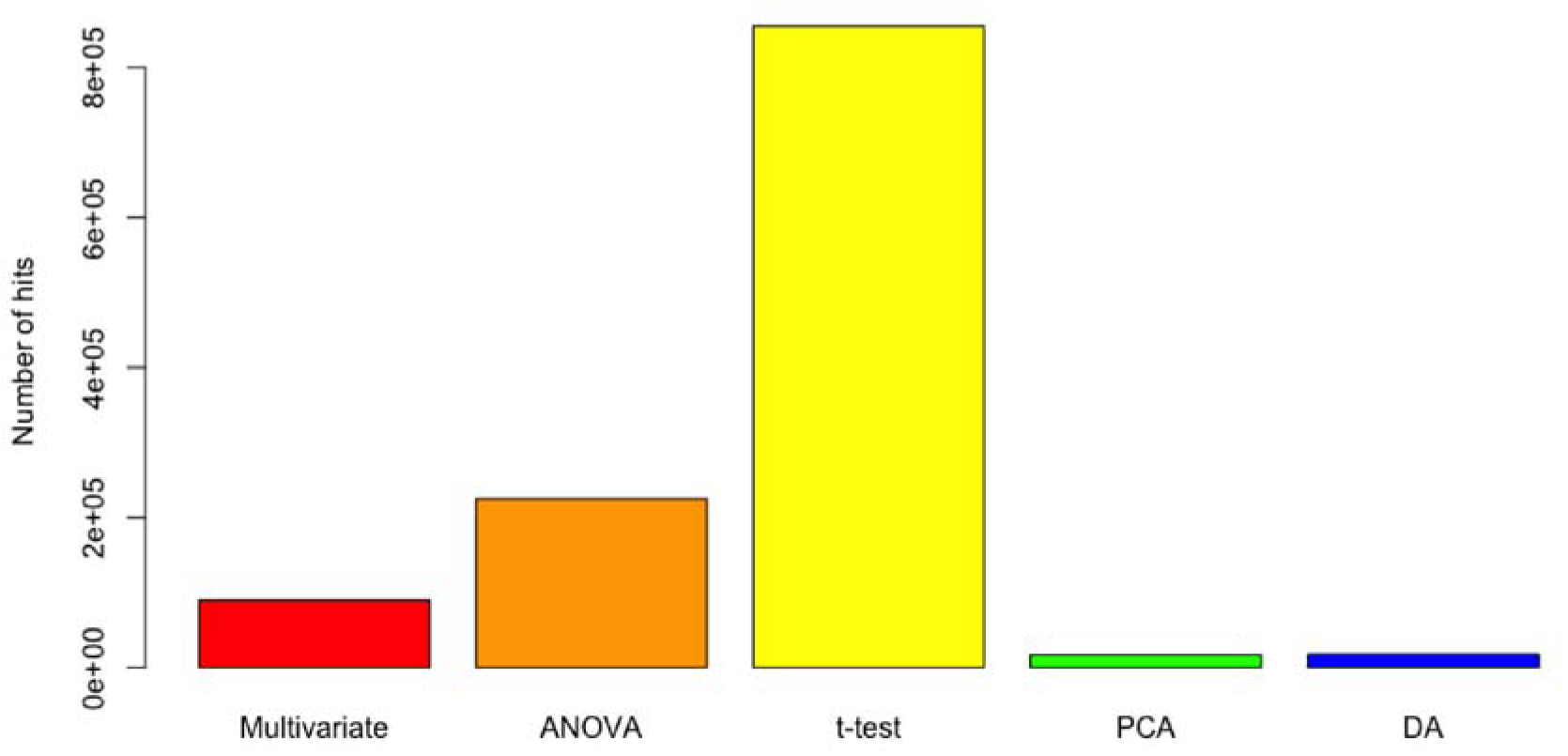
Popularity of analyses methods in Physiology. The number of hits from Google scholar using the search terms “Physiology AND multivariate”, “Physiology AND ANOVA”, “Physiology AND t-test”, “Physiology AND “principal component”” or “Physiology AND “discriminant analysis””. Note that Google Scholar was used since, unlike Web of Science or Pubmed it includes full text results.

Usage of principle component analyses (PCA), K-means clustering and other multivariate techniques have all been proposed previously for physiological analyses^1^, but still they are not used widely. Perhaps because they are thought of as too descriptive where specific hypothesis testing is required.

In this short report we compare the effectiveness of an established multivariate test (PCA) with a relatively new tool discriminant analyses of principal components (DAPC) from the package Adegenet^2^.

## Methods

The data analyses here are based on simulated experiments where agonist activates release of a second messenger throughout the cell. Two different agonists were used at a time, and we tested the hypothesis they both have the same effect on second messenger concentrations and distribution. The simulation was created using Python with the NEURON reaction diffusion (RXD) package^3^ embedded. Agonist is triggered to activate an enzyme *producer* located to 15 different membrane loci and the reaction produces the second messenger “*tfact*” which is in turn broken down by two different degrative enzymes, located either near the membrane or throughout cytosolic regions of the cell. Different agonists were assumed to evoke different producer enzyme *rates*. The enzyme rates were subject to 15% random Gaussian variation between experiments. Figure 2 shows example snapshots of the output data. Analyses were conducted using open source packages in R, FactoMineR^4^ for PCA, Adegenet for DAPC^2^ and ClusterSignificance^5^ to test the statistical significance of cluster separation. Median *p-value* (and confidence interval) for correct posterior classification of control and test following DAPC were calculated with the native Wilcox test in R.

**Figure 2:**
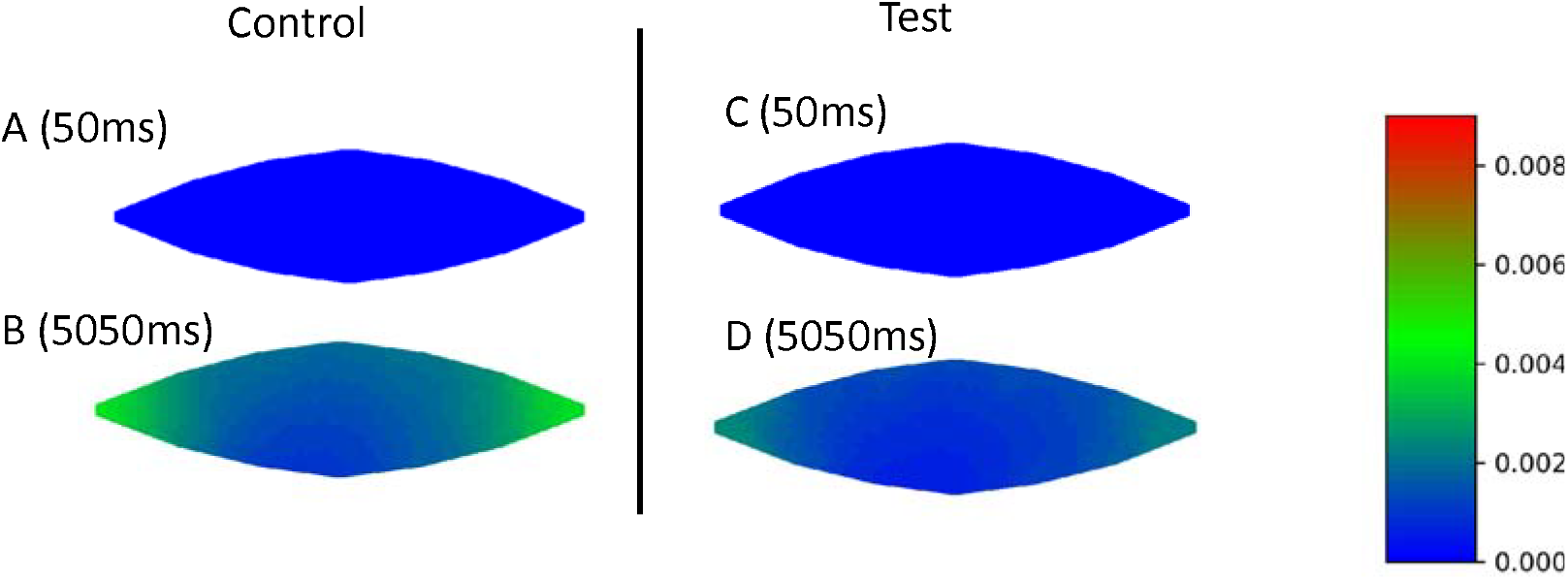
Snapshot of experiment output. The RXD (reaction diffusion model) output of tfact produced by a producer enzyme. **(A and B)** are control; **(A)** is just prior to the application of agonist (50ms), **(B)** is after 5 seconds of agonist (5.05s). **(C and D)** are test, where agonist potency has been halved. (C) is just prior to the application of agonist (50ms), **(D)** is after 5 seconds of agonist (5.05s).

## Results

We tested the ability of both PCA and DAPC to accurately detect differences in concentrations of *tfact* resulting from different simulated producer rates. We measured this over a spatial grid sliced through the centre of the cell (z=0). Figure 3 shows a snapshot of comparisons between *producer* rates of 2e-4 *mMms*^*-1*^ (control), and 1e-4 *mMms*^*-1*^ (test). With this arrangement and 20 (10, 10) simulated control and test experiments both PCA and DAPC performed well (Figure 3, 4). Cluster separation was significantly different by PCA, mean Euclidean separation distance was 0.09 arbitrary units (au) (95% CI: 0.084 to 0.089 au, *n*=10,10, *p-val*=2e-4).

**Figure 3:**
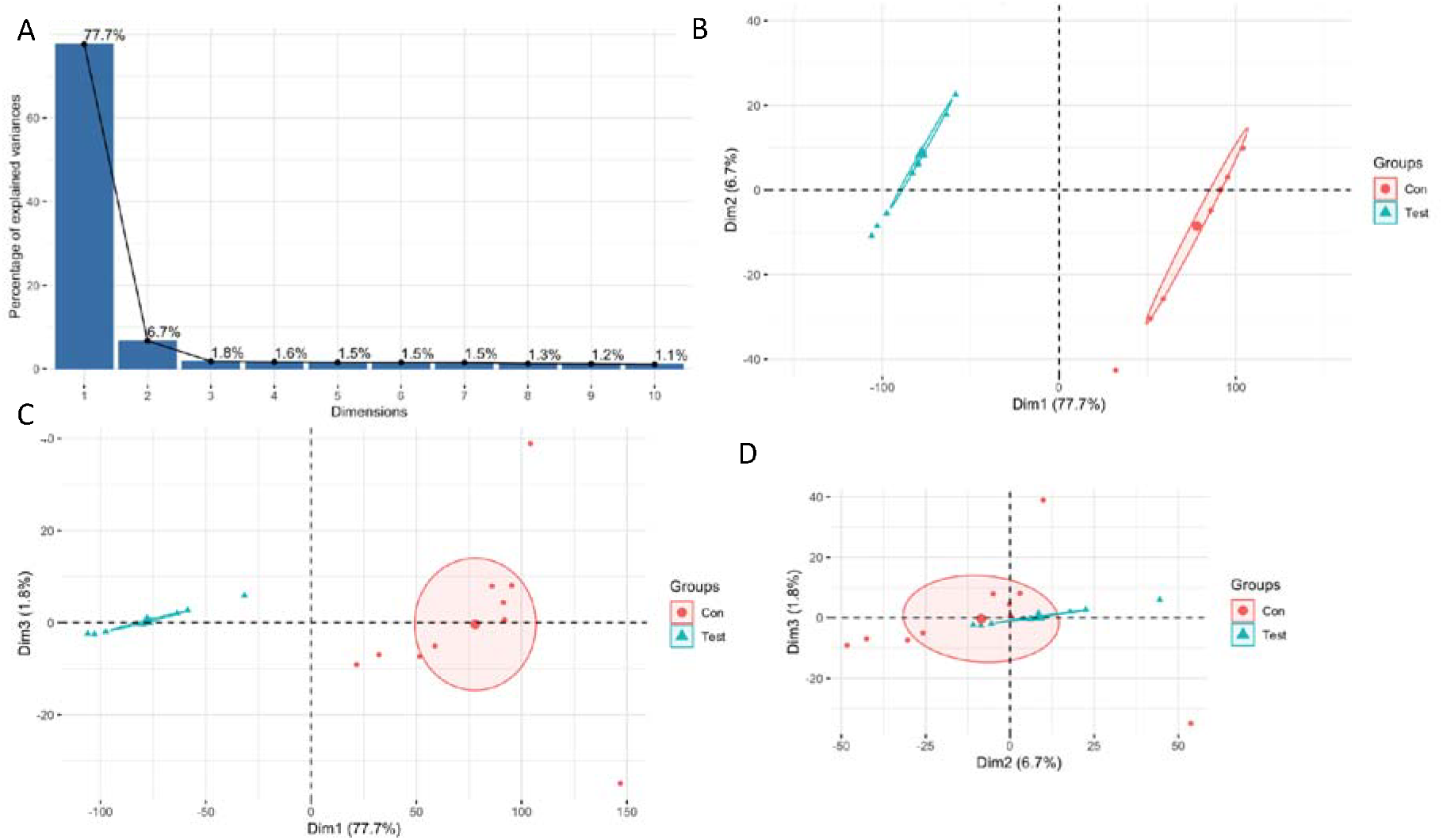
PCA analyses for halving agonist potency. **(A)** The Scree plot for PCA of the RXD dataset. **(B,C,D)** Cluster plots for the PCA, 1^st^ dimension/principal component verses 2^nd^ dimension (Dim1 vs Dim2), Dim 1 vs Dim 3, and Dim 2 vs Dim 3.

**Figure 4:**
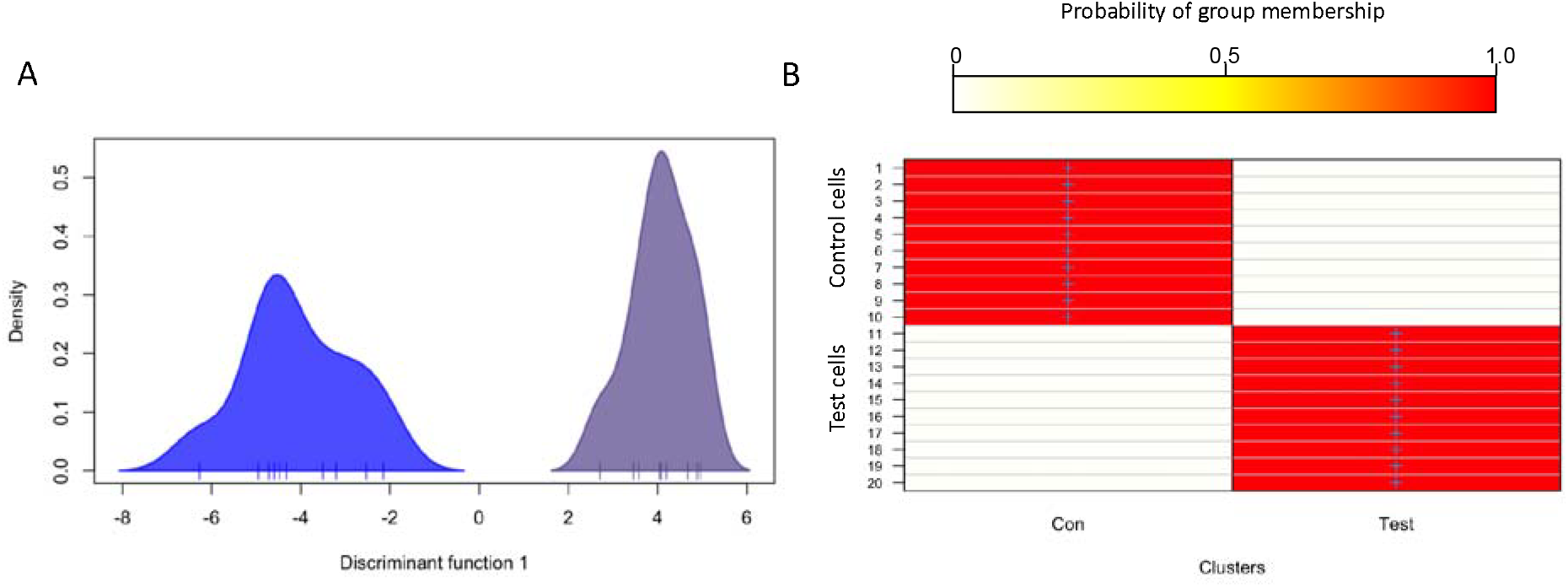
DAPC analyses for halving agonist potency. **(A)** Kernel plot for the 1^st^ (and only since there are only two groups) discriminant function (input 10 principal components), of the RXD dataset. **(B)** Posterior assignment heatmap. The scale bar above indicates the colour graded probabilities. The plot shows that each of the 10 control and 10 test (50% production rate) cells have been correctly identified, given a prior probability of 0.5. n=10 control and 10 test experiments.

For DAPC, and with 10 principal components (PC) input to the one discriminant function (one discriminant function because there are only two groups and thus one discrimination), there is a large and statistically significant separation, with posterior *p-value* for separation of groups averaging 0 (95% CI: 0 to 0), i.e., <1e-100 (n=10, 10, the same experiments as above). We then repeated this test with progressively smaller differences between the agonist potencies (Figure 5).

**Figure 5:**
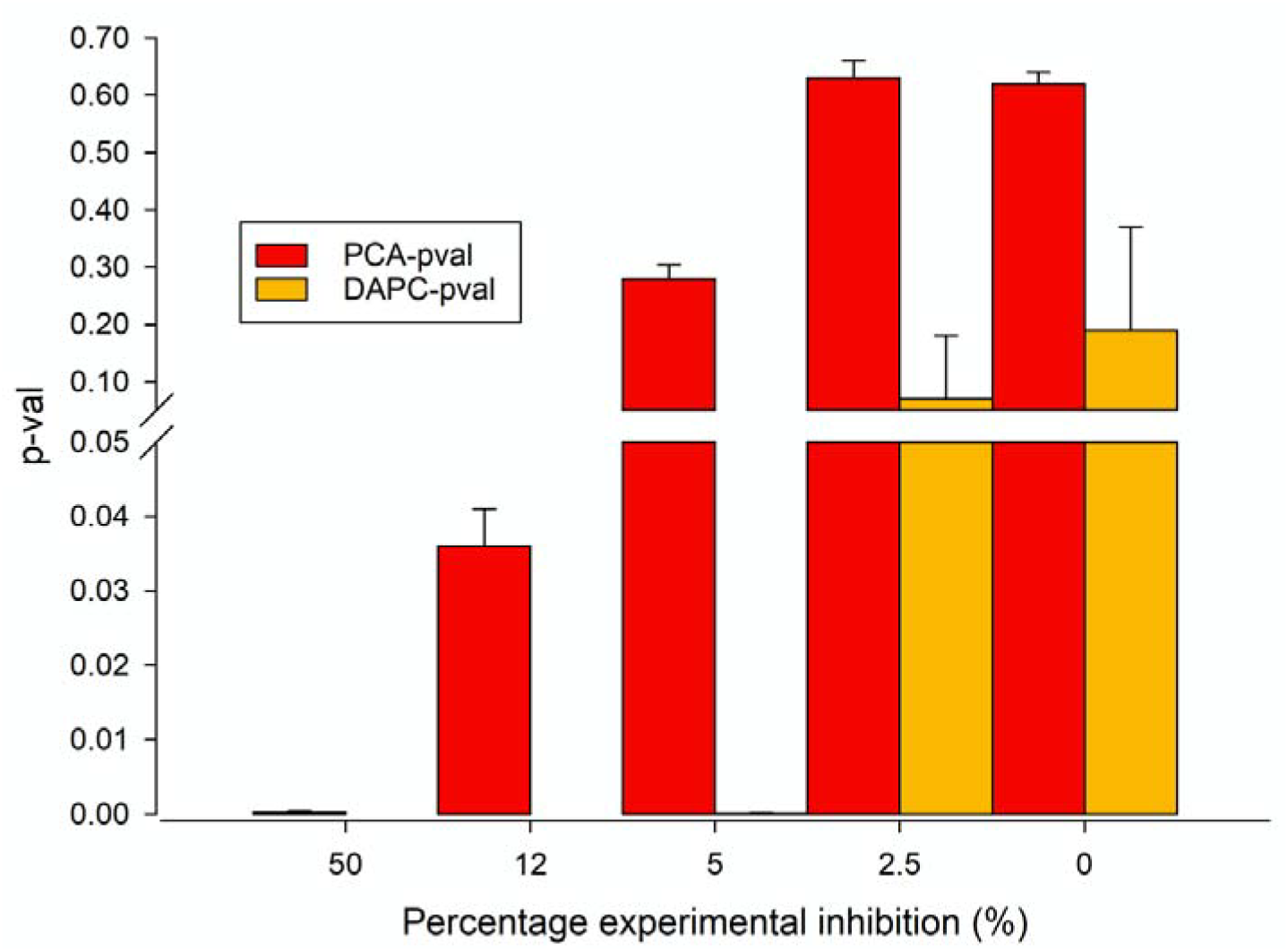
Snapshot of experiment output. Summary of the efficiency of PCA and DAPC to identify a significant effect of agonist change over the time and space of the experimental cells. The x-axis shows the percentage change in agonist potency; 50% mean rate decrease from 2e-4 mMms^−1^ to 1e-4, and 2.5% means a change from 2e-4 to 1.95e-4 mMms^−1^. 0 indicates a negative control where the two datasets were generated with the same rate equation albeit subject to gaussian random variation. N=10 control and 10 test, per experiment = 100 experiments total.

With 5% reduction of agonist potency (i.e., enzyme rate dropped from 2e-4 *mMms*^*-1*^ to 1.9e-4 *mMms*^*-1*^) the difference was no longer apparent with PCA (mean *p-value* for cluster separation, 0.28, 95% CI: 0.26 to0.313), but DAPC still separated these populations, with median posterior *p-value* of 1.69e-6 CI:5e-8 to 9e-5. As a negative control we simulated datasets with the same mean rates, and under these conditions, neither PCA or DAPC gave significant separation.

## Discussion/Conclusion

We have shown that the multivariate technique DAPC works well in the context of simulated real-time physiological experiments, detecting significance with much smaller physiological challenge than that needed for PCA. Furthermore, these analyses were conducted on image data across the entire dataset, although we constrained to just analyse the z=0 plane through the centre of the cell. There is a note of caution with the DAPC however. A critical parameter is the number of PCs chosen for the analyses. The more PCs chosen the better the separation between groups, but with enough PCs the algorithms would be powerful enough to create false positives. This potential problem of this over fitting can be controlled for with rigorous use of negative controls.

